# Cell size modulates ferroptosis susceptibility

**DOI:** 10.1101/2025.10.25.684524

**Authors:** Evgeny Zatulovskiy, Magdalena B. Murray, Shuyuan Zhang, Scott J. Dixon, Jan M. Skotheim

**Author notes:** Correspondence (E.Z.), (J.M.S.).

## Abstract

Size is a fundamental property of cells that influences many aspects of their physiology. This is because cell size sets the scale for all subcellular components and drives changes in the composition of the proteome. Given that large and small cells differ in their biochemical composition, we hypothesize that they should also differ in how they respond to signals and make decisions. Here, we investigated how cell size affects susceptibility to cell death. We found that large cells are more resistant to ferroptosis caused by system x_c_^-^ inhibition. Ferroptosis is a type of cell death characterized by the iron-dependent accumulation of toxic lipid peroxides. This process is opposed by cysteine-dependent lipid peroxide detoxification mechanisms. We found that larger cells exhibit higher concentrations of the cysteine-containing metabolite glutathione and lower concentrations of membrane lipid peroxides. Mechanistically, this can be explained by the fact that larger cells had lower concentrations of an enzyme that enriches cellular membranes with peroxidation-prone polyunsaturated fatty acids, ACSL4, and increased concentrations of the iron-chelating protein ferritin, the glutathione-producing enzymes glutamate-cysteine ligase and glutathione synthetase, and the lysosomal protease cathepsin B, which can catabolize cysteine-rich extracellular proteins to produce additional cystine for fueling the synthesis of glutathione. Taken together, our results highlight the significant impact of cell size on cellular function and survival, revealing a size-dependent vulnerability to ferroptosis that could influence therapeutic strategies based on this cell death pathway.

## INTRODUCTION

Cell size is fundamental to cell physiology. This is because it impacts cell geometry and sets the scale of all biosynthetic processes in the cell (Chan & Marshall, 2010; Ginzberg et al, 2015; Zatulovskiy & Skotheim, 2020). The importance of cell size is most clearly seen when yeast and mammalian cells become excessively large. In these cases, cells are not able to maintain optimal protein and mRNA concentrations to support growth as their cytoplasm becomes more dilute, unless the increase in cell size is coulter-balanced by the increase in cell ploidy (Neurohr et al, 2019; Lanz et al, 2022, 2024; Zatulovskiy et al, 2022). Such excessively large cells exhibit features of senescence and are no longer able to enter the cell division cycle. Unsurprisingly, cell size regulation is compromised in many diseases and during aging (Sandlin et al, 2022; Nguyen et al, 2016; Lengefeld et al, 2021). As animals, including humans, grow older, they contain more ‘senescent’ cells that tend to be larger (Davies et al, 2022). For example, basal keratinocytes are about 1.5 times as large when humans are 80 years old versus 20 (Liao et al, 2012). Conversely, cancer cells often exhibit an extreme heterogeneity in cell size, which may be associated with more proliferation (Sandlin et al, 2022; Nguyen et al, 2016; Li et al, 2015; Bell & Waizbard, 1986). Taken together, these studies argue for the importance of cell size in both natural and disease contexts.

Due to the importance of cell size for cell physiology, this property is highly regulated by distinct classes of molecular mechanisms. One class of mechanisms regulates the cell growth rate so that smaller cells grow faster and add proportionally more biomass than larger cells within one cell division cycle to ensure that all cells divide at similar sizes (Ginzberg et al, 2018; Liu et al, 2024; Conlon & Raff, 2003). A second class of molecular mechanisms, useful in more regular shaped cell types like rod shaped bacteria and fission yeast, relies on geometrical parameters such as cell length or surface area to measure cell size and determine that appropriate timing for initiating cell division (Miller et al, 2023; Martin & Berthelot-Grosjean, 2009; Pan et al, 2014; Bhatia et al, 2014; Moseley & Nurse, 2010). Finally, a third class of mechanisms rely on differential changes in protein concentrations of cell cycle activators and cell cycle inhibitors as cells grow larger (Chen et al, 2020b; Zatulovskiy et al, 2020; Schmoller et al, 2015). Typically, cell cycle activators either remain at constant concentration or increase with cell size, while some key cell cycle inhibitors, like the transcriptional inhibitors Whi5 in budding yeast and the retinoblastoma protein RB in human cells, are progressively diluted in larger cells to trigger their division (Zatulovskiy et al, 2020; Schmoller et al, 2015; Zhang et al, 2022b, 2024).

That some larger cells initiate division through increases in the concentrations of cell cycle activators relative to cell cycle inhibitors was surprising because previous bulk studies showed that total protein concentration remained constant as cells grew larger. However, this constant concentration of total protein masked changes in the concentrations of individual proteins as cells grow larger, as revealed by a systematic quantitative proteomics analysis (Lanz et al, 2022, 2024; Zatulovskiy et al, 2022). The proteomics analyses revealed that as cell size increases, the concentrations of numerous senescence-related proteins begin to approach those of a senescent cell, even before cells lose the ability to enter the cell division cycle (Lanz et al, 2022; Crozier et al, 2023). This may in part explain the association of large cell size with the senescent state. Moreover, large cells have difficulty in replicating and repairing their genomes, which may be related to the lower concentrations of DNA replication and repair enzymes observed in larger cells (Lanz et al, 2022; Crozier et al, 2022; Manohar et al, 2023; Foy et al, 2023; Wilson et al, 2023). Taken together, these results indicate that a cell size increase in the absence of DNA replication impacts cell physiology through changing the composition of the proteome.

If cell size drives widespread changes in proteome composition, then this might impact many areas of cell physiology, including the susceptibility to cell death. Indeed, proteomic and transcriptomic comparisons of different-sized cells within the same cell type identified concentration changes for many genes involved in regulated cell death (Lanz et al, 2022; Miettinen & Björklund, 2016). Consistent with this, cell size is correlated with some cell death decisions – smaller cells have a higher likelihood to undergo apoptosis both in cell cultures and during *C. elegans* and *D. melanogaster* development (Miettinen & Björklund, 2016; Sethi et al, 2022; Kiyomitsu & Cheeseman, 2013; Hatzold & Conradt, 2008; Chen et al, 2016; Cordes et al, 2006). Inspired by these studies, we aimed to mechanistically investigate how cell size affects different cell death responses in human cells, as it remains unknown whether cell size impacts non-apoptotic cell death.

To test if there is a relationship between cell size and cell death, we used a high-throughput microscopy based approach to examine a variety of stresses (Forcina et al, 2017; Inde et al, 2020). We found that susceptibility to ferroptosis, an iron-dependent form of cell death, demonstrated the strongest dependence on cell size. This finding resonates with a recent report that cell susceptibility to RSL3, a chemical that induces ferroptosis by inhibiting the reduction of lipid peroxides by glutathione peroxidase 4 (GPX4), increases with cell size (Chan et al, 2025). Inhibition of the cystine/glutamate antiporter system x_c_^-^, using the small molecule erastin2 (Era2) (Dixon et al, 2014) preferentially induced cell death in smaller cells, while larger cells were more resistant. Ferroptosis is characterized by the accumulation of toxic lipid peroxides (Jiang et al, 2021; Dixon & Olzmann, 2024). To suppress this accumulation and prevent ferroptosis, cells can use cystine/cysteine-derived metabolites, like the reduced tripeptide glutathione (GSH), to prevent the accumulation of peroxidized lipids. The lower susceptibility of large cells to Era2-induced ferroptosis can likely be explained by our observation that larger cells are able to produce higher amounts of GSH per unit protein mass, and that they have a lower concentration of enzymes like ACSL4 that incorporate oxidizable polyunsaturated fatty acids into the plasma membrane (Doll et al, 2017). Supporting this model, the genetic disruption of *ACSL4* reduced the size dependence of lipid peroxidation and ferroptotic cell death. Taken together, our results show how large cell size can protect against ferroptosis.

## RESULTS

### Larger cells are less susceptible to ferroptosis

To test our hypothesis that cell size affects susceptibility to cell death, we exposed cells of different sizes to a variety of compounds known to induce different forms of cell death. To do this, we first used fluorescence-activated cell sorting (FACS) to isolate small and large G1-phase HMEC*-hTERT* cells (human mammary epithelium cells immortalized with telomerase, herein referred to as HMEC for brevity) (Fig. 1A,B) (Lanz et al, 2022). We sorted cells for size using side scatter, which correlates well with cell size (Lanz et al, 2022; Tzur et al, 2011; Berenson et al, 2019). In this experiment we used DNA staining with Hoechst dye to only collect the cells that were in G1 phase of the cell cycle, since the cell cycle phase can influence cell survival (Rodencal et al, 2024; Ruiz-Losada et al, 2022; Lee et al, 2024; Kuganesan et al, 2023). The small- and large-sorted cells were then seeded into 384-well plates and treated with death-inducing compounds over a range of different doses. Each population of cells was then monitored using high-throughput fluorescence microscopy for 72 h to measure dose-dependent responses and cell death kinetics (Forcina et al, 2017; Inde et al, 2020).

**Figure 1.**
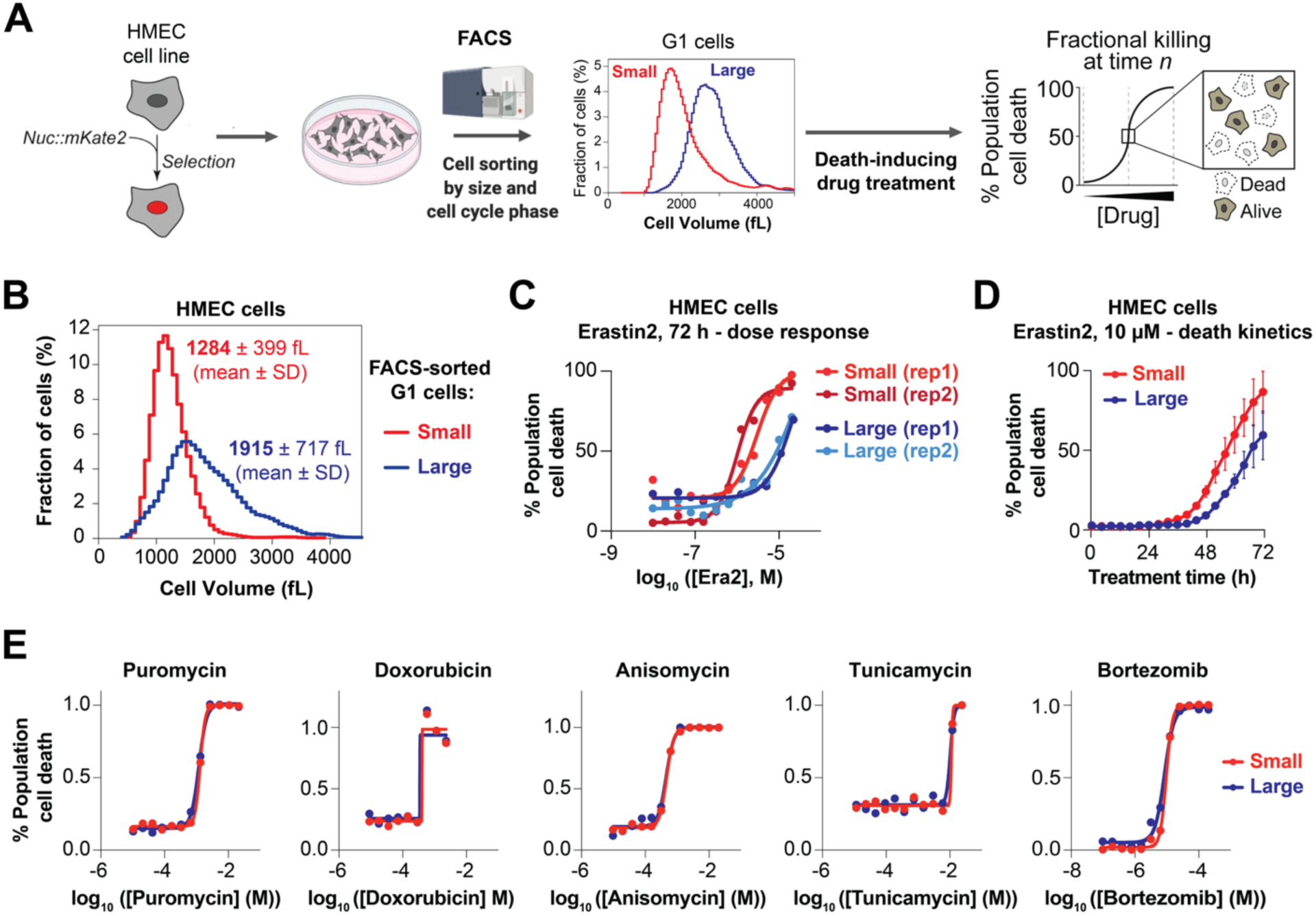
High-throughput microscopy-based measurement of cell death susceptibility in different-sized cells. (A) Experiment schematic: a genetic construct encoding nuclear-localized fluorescent protein mKate2 was delivered into HMEC cells using lentiviral transduction. After selection, these cells were FACS-sorted by cell cycle phase and cell size into small and large G1 cells (smallest and largest 5%). The FACS-sorted cells were seeded on 384-well plates and allowed to settle overnight before being treated with death-inducing compounds in the media containing a dead-cell dye SYTOX Green. The cells were imaged for 72 h during treatment, and live cells were identified by the presence of nuclear mKate2 fluorescence, while dead cells were identified by the presence of the SYTOX Green signal (Forcina et al, 2017; Inde et al, 2020). The numbers of live and dead cells were automatically tracked over time to determine the dose-response curves and cell death kinetics. (B) Cell size distributions of small and large G1-phase HMEC cells after FACS sorting, measured on a Coulter counter. (C) Dose-response curves of small and large FACS-sorted HMEC cells treated with a potent ferroptosis-inducing compound erastin2 (Era2). Rep1 and rep2 denote two different biological replicates, each with two technical replicates. (D) Cell death kinetics of small and large FACS-sorted HMEC cells treated with 10 µM Era2. Dots show the means of two biological replicates, error bars denote the range. (E) Dose-response curves of small and large FACS-sorted HMEC cells treated for 72 h with other lethal compounds: puromycin, doxorubicin, anisomycin, tunicamycin and bortezomib. Each experiment was performed twice independently, and each biological replicate included two technical replicates. The dots show averages of two biological replicates.

Comparing dose-response curves for small and large cells across different compounds, we observed the largest size-dependent differences in cell death susceptibility in response to erastin2 (Era2). Era2 induces ferroptosis by inhibiting system x_c_^-^ (Dixon et al, 2014). Larger HMEC cells were resistant to higher doses of Era2 than smaller cells. For example, at the 72 h time point, the Era2 IC_50_ was 28 ± 11 µM (mean ± SD) for large cells versus 2.0 ± 1.4 µM for small cells (Student’s t-test: *p = 0.039*) (Fig. 1C). Moreover, larger cells exhibit slower cell death kinetics (Fig. 1D). By contrast, for compounds that cause cell death by disrupting protein synthesis (puromycin, anisomycin), inducing DNA damage (doxorubicin), inhibiting protein glycosylation (tunicamycin), or blocking proteasome function (bortezomib), IC_50_ values were not statistically significantly different for large versus small cells (Fig. 1E). We note that this does not eliminate the possibility that there are size-dependent differences in susceptibilities to these compounds that would manifest if our size range were expanded further. However, since the most pronounced differences were observed for Era2, we focused our investigation on how cell size affects ferroptosis.

To test if protection against ferroptosis by large cell size was generalizable beyond HMEC cells isolated by FACS, we used alternative methods to generate different sized populations of cells and tested additional cell lines. We examined the HT-1080 fibrosarcoma cell line, commonly used in ferroptosis studies (Dixon et al, 2012), and non-transformed telomerase-immortalized retinal pigment epithelium cells (RPE-1 cell line), commonly used in cell cycle and cell size studies. We used FACS to sort G1-phase HT-1080 cells into four cell size bins and treated these cells with 0.3 µM Era2. As observed in HMEC cells, larger HT-1080 and RPE-1 cells were more resistant to ferroptotic cell death (Fig. S1A). Next, to generate populations of different-sized cells without cell sorting, we arrested HT-1080 cells in early G1 using the cell-cycle inhibitor palbociclib (CDK4/6 inhibitor) for 2-6 days (Neurohr et al, 2019; Lanz et al, 2022; Crozier et al, 2023; Manohar et al, 2023; Foy et al, 2023). Larger cells, which were arrested in G1 for a longer time, were less sensitive to Era2 than smaller cells arrested for a shorter duration (Fig. 2A,B). We note that non-arrested cells had a lower susceptibility to Era2-induced ferroptosis compared to cells that were arrested in G1 for 2-3 days, despite being smaller in size. This is likely due to the difference in the fraction of cells in different cell cycle phases between arrested and non-arrested conditions since cells in S/G2/M phases are known to be more resistant to ferroptosis than cells in G0/G1 phases (Rodencal et al, 2024; Kuganesan et al, 2023).

**Figure 2.**
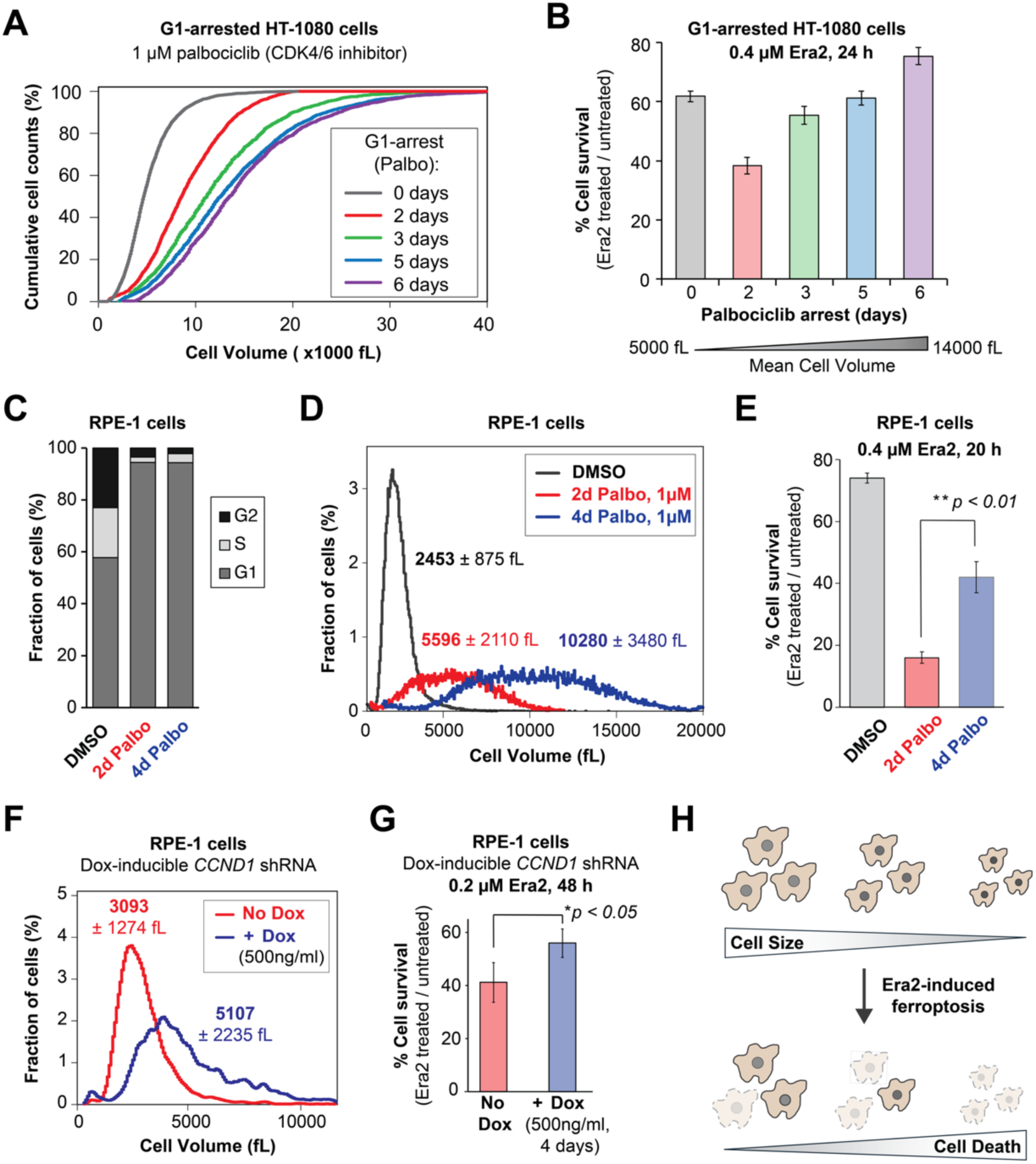
Larger cells are less susceptible to Era2-induced ferroptosis. (A) Cumulative cell size distributions of HT-1080 cells, whose cell cycle was arrested in G1 phase for 0, 2, 3, 5, or 6 days with 1 µM of the CDK4/6 inhibitor palbociclib. (B) Cell survival percentage of HT-1080 cells whose size was increased through palbociclib-induced cell cycle arrest. After pre-treatment with palbociclib for the indicated numbers of days, the cells were exposed to 0.4 µM Era2 in the presence of palbociclib for 24 h. Cell survival percentage was calculated relative to the cells treated only with palbociclib. (C) Cell cycle phase distribution of asynchronously proliferating RPE-1 cells, and RPE-1 cells treated with the CDK4/6 inhibitor palbociclib for 2 or 4 days. (D) Cell size distribution of asynchronously proliferating RPE-1 cells, and RPE-1 cells treated with the CDK4/6 inhibitor palbociclib for 2 or 4 days. The numbers next to the histograms indicate mean cell size ± standard deviation for the corresponding condition. (E) Cell survival percentage in RPE-1 cells whose cell cycle was synchronized and size was increased through 2-day or 4-day palbociclib-induced cell cycle arrest. After pre-treatment with palbociclib for the indicated numbers of days, the cells were exposed to 0.4 µM Era2 in the presence of palbociclib for 20 h. Cell survival percentage was calculated relative to the cells treated only with palbociclib. (F) Cell size distributions of RPE-1 cells that inducibly express shRNA against cyclin D1 gene (*CCND1*) to slow down the cell cycle and increase cell size. Cells were grown in medium containing 500 ng/mL doxycycline for 4 days to induce shRNA expression and increase cell size. Uninduced cells were grown in the same medium with DMSO in place of doxycycline. (G) Cell survival percentage in control RPE-1 cells and cells expressing an shRNA against cyclin D1 gene (*CCND1*). Cells were treated with 0.2 µM Era2 for 48 h, and cell survival percentage was calculated relative to DMSO-treated cells. Cell survival percentages in graphs (B), (E) and (G) are shown as means ± s.e.m.; n = 3 biological replicates. A two-tailed unpaired Student’s t-test was used to evaluate the statistical significance of survival percentage differences in panels (E) and (G). (H) Graphical summary: Larger cells are less susceptible to Era2-induced ferroptosis.

We further confirmed that the effects of cell size on Era2-induced ferroptosis susceptibility are generalizable across cell lines and independent of the cell cycle. To do this, we treated RPE1 cells with palbociclib for 2 or 4 days and then measured their susceptibility to Era2. Both the cells treated with palbociclib for 2 days and for 4 days were arrested in G1, but the cells treated for 4 days were nearly two-fold larger than the cells treated for 2 days (Fig 2C,D). This enabled us to isolate the effects of cell size from cell cycle effects and confirm that larger cells were significantly more resistant to Era2-induced ferroptosis compared to their smaller counterparts in the same phase of the cell cycle (Fig 2E).

As an additional means to generate populations of different sized cells, we used inducible shRNA-mediated cyclin D1 (*CCND1*) knockdown in RPE-1 cells to generate a population of cells that had larger cell sizes compared to uninduced RPE-1 cells (You et al, 2025). As previously reported, cells that inducibly expressed *CCND1* shRNA had a ∼30% decrease in Cyclin D1 protein concentration but continued to grow and proliferate, though at a lower division rate (You et al, 2025). As a result, these *CCND1* knockdown cells were larger than uninduced cells, and were less susceptible to Era2 (Fig. 2F,G).

Our finding that larger cells are less susceptible to ferroptosis induced by Era2 treatment seemingly contradicted findings that large size sensitizes cells to the ferroptosis-inducing compound RSL3 (Chan et al, 2025). Unlike Era2, which acts by inhibiting cystine/glutamate antiporter system x_c_^-^ and thereby depleting glutathione (GSH) pool, RSL3 belongs to a different class of ferroptosis inducers, which acts by inhibiting glutathione peroxidase 4 (GPX4) (Yang et al, 2014), a key enzyme that uses GSH to reduce toxic lipid peroxidation. To compare the effects of these two classes of compounds in our experimental system, we treated the HT-1080 cells, arrested in G1 for 2-6 days using palbociclib, with RSL3 (Fig. S1B). Consistent with the published report (Chan et al, 2025) and in contrast to Era2 treatment (Fig. 2B), increased cell size during G1 arrest progressively increased sensitivity to RSL3. This difference in cell responses to the two different classes of ferroptosis inducers has been noted in several reports describing context-dependent responses to different classes of ferroptosis inducers (Chan et al, 2025; Rodencal et al, 2024; Magtanong et al, 2022).

Taken together, our measurements demonstrate that in human cell cultures, larger cells are generally less susceptible to ferroptosis induced by system x_c_^-^ inhibition (Fig. 2H). We observed this result in populations of different sized cells that were generated through multiple methods including genetics, FACS sorting, and small molecule-induced cell cycle arrest.

### Membrane lipid peroxidation decreases with cell size

Ferroptosis is defined and caused by the accumulation of toxic products from polyunsaturated fatty acid (PUFA) peroxidation in the plasma membrane (Jiang et al, 2021; Dixon & Olzmann, 2024) (Fig. 3A). We therefore next sought to determine how cell size affects ferroptosis-associated lipid peroxidation. Lipid peroxidation can be detected using a ratiometric fluorescent sensor dye BODIPY-C11 581/591 (Dixon et al, 2012; Pap et al, 1999). This dye can be used for ratiometric analysis of lipid peroxidation in flow cytometry, which detects a shift of the fluorescence emission peak from red (∼590 nm) to green (∼510 nm) caused by the oxidation of the polyunsaturated butadienyl portion of this fatty acid analog. When HMEC cells are treated with Era2, which leads to increased lipid peroxidation, we find a decrease in non-oxidized (red) BODIPY-C11 fluorescence and an increase in oxidized (green) BODIPY-C11 as expected (Fig. 3B).

**Figure 3.**
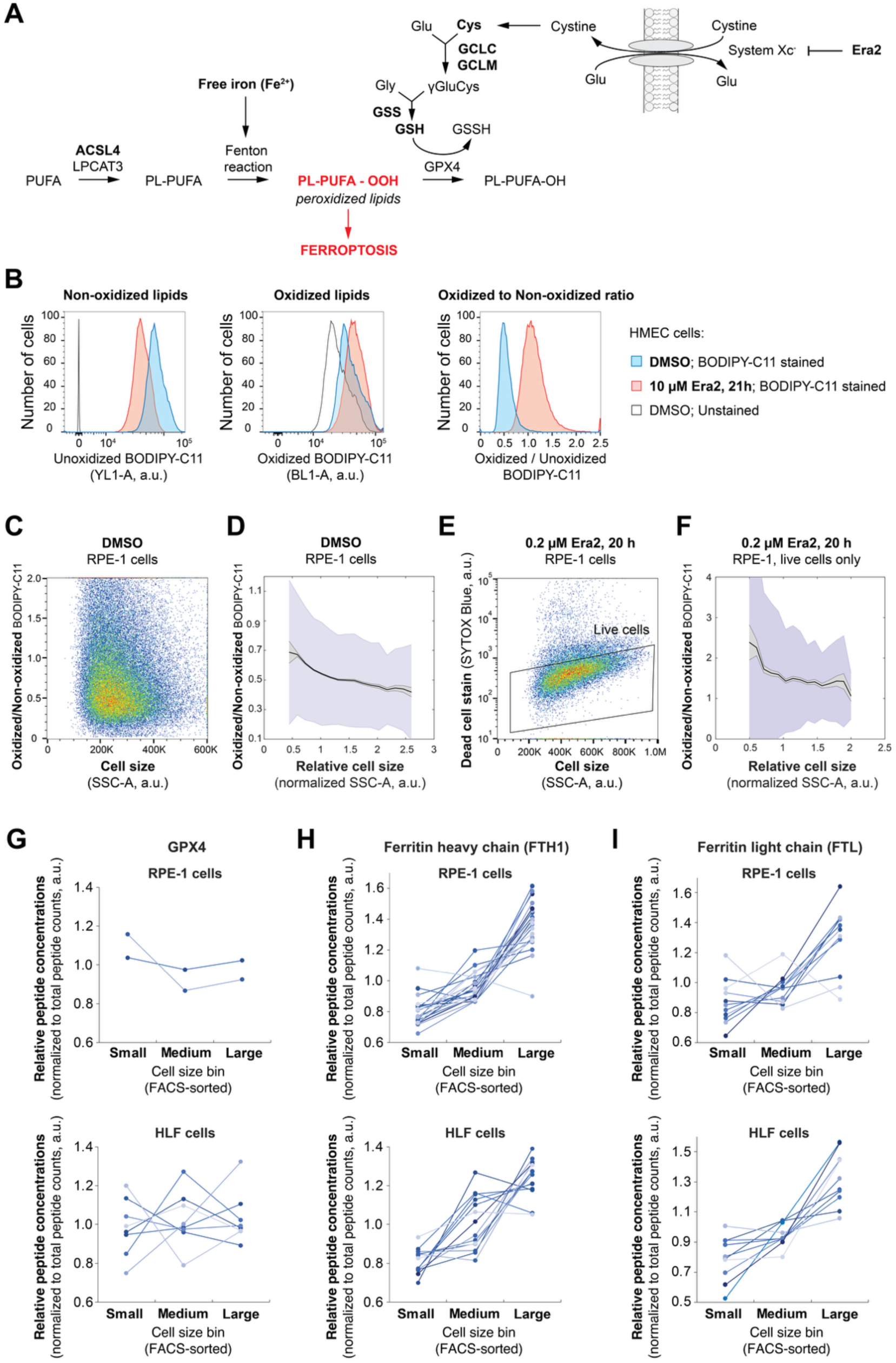
Membrane lipid peroxidation decreases with cell size. (A) Schematic of pathways regulating ferroptosis. Ferroptosis is induced through the accumulation of toxic peroxidized lipid species in the plasma membrane. The accumulation of peroxidized lipids is prevented by glutathione (GSH) production and GSH-dependent reduction of the peroxidized lipids (Jiang et al, 2021; Dixon & Olzmann, 2024). (B) The ratiometric fluorescent dye BODIPY-C11 581/591 detects lipid peroxidation in live HMEC cells (Dixon et al, 2012; Pap et al, 1999). Oxidation of the polyunsaturated butadienyl portion of this fatty acid analog results in a shift of the fluorescence emission peak from red (∼590 nm) to green (∼510 nm), allowing ratiometric analysis of lipid peroxidation using flow cytometry. Treatment of cells with Era2 increases lipid peroxidation, as indicated by decreased non-oxidized (red) BODIPY-C11 fluorescence and increased oxidized (green) BODIPY-C11 fluorescence. (C) A flow cytometry analysis of lipid peroxidation in different-sized RPE-1 cells within the DMSO-treated cell culture. The side-scatter parameter (SSC-A) was used as a proxy for cell size (Lanz et al, 2022; Tzur et al, 2011). Three biological replicates were performed, and 100,000 events were recorded for each. (D) A quantitative analysis of the flow cytometry data from panel (C): the data were binned into 12 bins by normalized cell size (SSC-A / median SSC-A), and the mean values for of oxidized (green) to non-oxidized (red) BODIPY-C11 ratio were plotted for each bin (black line). Gray shaded area denotes the s.e.m. for each bin, and the blue area denotes the standard deviation. (E) For Era2-treated cell populations, live cells were identified and gated in the indicated region using a live/dead cell permeability dye SYTOX Blue. (F) Size-dependent lipid peroxidation in RPE-1 cells treated with 0.2 µM Era2 for 20 h. The analysis was performed as in panel (D). (G-I) Proteomics-based analysis of GPX4 (G), ferritin heavy chain (H) and ferritin light chain (I) expression in FACS-sorted small, medium and large RPE-1 cells and primary lung fibroblasts (HLF). Each line in the plot corresponds to a unique peptide from the indicated proteins identified by mass spectrometry reported in *Lanz et al.* (Lanz et al, 2022).

To determine how cell size affects basal lipid peroxidation in cells, we first stained control (DMSO-treated) cells with BODIPY-C11 and subjected them to a flow cytometry analysis using side scatter (SSC-A) as a proxy for cell size (Lanz et al, 2022; Tzur et al, 2011). When we plotted the oxidized to non-oxidized BODIPY-C11 ratio against cell size, we observed a clear anti-correlation between these two parameters (Fig. 3C,D). We then performed a similar analysis on Era2-treated cells. For this analysis, we gated only live cells using a fluorescent cell permeability dye SYTOX Blue that stains dead or dying cells to omit cells undergoing Era2-induced death (Fig. 3E). This gating allowed us to exclusively focus on the live cells that were not dying. We found that in Era2-treated cell populations, larger cells demonstrated lower levels of lipid peroxidation, similarly to untreated cells (Fig. 3F). By contrast, membrane lipid peroxidation, the terminal cause of ferroptosis, exhibits higher basal levels in smaller cells. This provided a rationale for why smaller cells would require less treatment with Era2 to accumulate enough lipid peroxidation and induce ferroptosis.

Having determined that membrane lipid peroxidation decreases with cell size, we next investigated how the expression of key cellular components regulating this process changes with cell size. The overall lipid oxidation status in the membrane is set largely by the competition between Fe^2+^-dependent peroxidation of PUFA-containing phospholipids, and glutathione-dependent reduction of peroxidized lipids by GPX4 (Fig. 3A). We analyzed size-dependent proteomics data for RPE-1 cells and primary human lung fibroblasts (HLF) and observed no significant size-differences in GPX4 expression (Fig. 3G). However, ferritin heavy chain (FTH1) and ferritin light chain (FTL) concentrations increased by 1.6-to-2-fold with a two-fold increase in cell size (Fig. 3H,I). Ferritin acts as a key iron-storage protein that can promote ferroptosis resistance through iron chelation (Zhang et al, 2022a; Chen et al, 2020a). The observed increase in ferritin concentration with cell size could therefore lead to additional Fe^2+^ ion chelation, which in turn would protect large cells from iron-dependent lipid peroxidation and ferroptosis. However, when we measured the concentration of labile intracellular Fe^2+^ using a fluorescent probe FerroOrange (Hirayama et al, 2020), we did not observe any size-dependent decrease in labile iron concentration (Fig. S2A). Previous work suggests a link between increased sequestration of ferrous iron in lysosomes and resistance to ferroptosis. It was reported that senescent cells, which are also large (Fig. S3A,B), gain resistance to ferroptosis through lysosomal alkalinization and sequestration of ferrous iron in lysosomes (Loo et al, 2025). We therefore tested whether the superscaling of lysosomes observed in large cells (Lanz et al, 2022; You et al, 2025) promotes Era2 resistance through lysosomal iron sequestration. To do this, we stained the cells with the lysosomal iron detection probe Lyso-FerroRed (Saimoto et al, 2025) and measured its scaling using flow cytometry (Fig. S2B). We observed that the amount of Lyso-FerroRed, and therefore, the amount of lysosomal iron, scaled in direct proportion to cell size, just like the total cellular protein content (Fig. S2B). These results indicate that iron chelation by ferritin and its sequestration in lysosomes are unlikely to play a crucial role in size-dependent decrease in Era2 sensitivity.

### Larger cells have higher concentrations of glutathione

Having shown that lipid peroxidation decreases with cell size and that such a decrease is unlikely to be mediated by changes in labile Fe^2+^ concentration, we next sought to test if cell size also affects glutathione (GSH) abundance, which contributes to oxidized lipid reduction and ferroptosis resistance, and is depleted in cells upon system x_c_^-^ inhibition (but not GPX4 inhibition) (Jiang et al, 2021; Dixon & Olzmann, 2024; Dixon et al, 2012) (Fig. 3A). To determine how cell size affects glutathione synthesis, we re-analyzed our previously collected proteomics data, where we compared the proteomes of FACS-sorted small, medium, and large RPE-1 cells (Lanz et al, 2022) (Fig. 3). We found that concentrations of two key proteins involved in GSH biosynthesis, glutamate-cysteine ligase modifier subunit (GCLM) and glutathione synthetase (GSS), increase with cell size. This indicates that larger cells might have higher GSH production rates (Fig. 4A).

**Figure 4.**
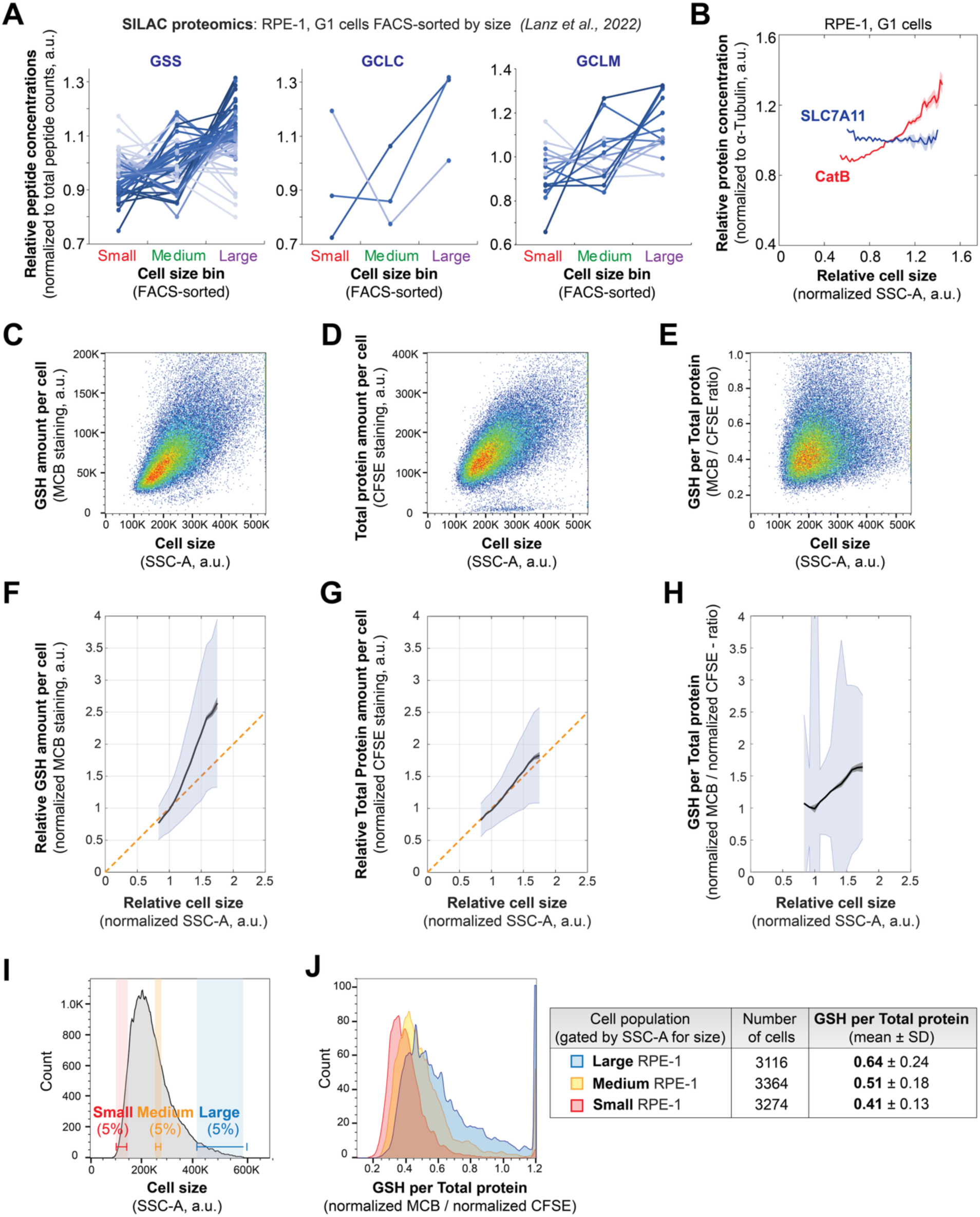
Larger cells have higher concentrations of glutathione and enzymes promoting glutathione synthesis. (A) Proteomics-based analysis indicates the concentrations of key enzymes involved in glutathione production (see Fig. 3A) increase with cell size. For this analysis, RPE-1 cells were FACS-sorted into populations of small, medium, and large G1 cells, and the proteomes of cells in these bins were analyzed using SILAC mass spectrometry. The primary proteomics data used to plot the concentrations of glutathione synthetase (GSS), glutamate-cysteine ligase catalytic (GCLC) and modifier (GCLM) subunits were taken from our previous work (Lanz et al, 2022). Each line in the plot corresponds to a unique peptide corresponding to the indicated protein that was identified by mass spectrometry. (B) Flow-cytometry-based measurement of cystine/glutamate transporter SLC7A11 (xCT) and cathepsin B (CatB) concentrations in G1-phase RPE-1 cells demonstrates a modest decrease in SLC7A11 and a significant increase in cathepsin B concentrations with cell size. To calculate the concentrations of SLC7A11 and CatB, their amounts were measured with flow cytometry using immunofluorescence and normalized to the amounts of α-Tubulin. The data were binned by cell size, and mean values for each bin were plotted against normalized cell size (solid blue line for SLC7A11 and red line for CatB). Shaded areas denote the s.e.m. for each bin. (C, D) Flow-cytometry-based measurement of GSH amount (C) and total protein amount (D) scaling with cell size in RPE-1 cells. The side scatter parameter (SSC-A) is used as a proxy for cell size (Tzur et al, 2011; Berenson et al, 2019). Three biological replicates were performed, and 100,000 events were recorded in each replicate. (E) GSH concentration in RPE-1 cells plotted against cell size. The GSH concentration is calculated as the ratio of GSH amount to the amount of total protein from data shown in panels (C) and (D). (F-H) Analysis of the flow cytometry data shown in panels (C-E). The data were binned by cell size, and mean values (black lines in the plots) for GSH amount (F), total protein amount (G) and GSH concentration (H) were plotted against normalized cell size (SSC-A / median SSC-A). Gray shaded areas denote the s.e.m. for each bin, and the blue area denotes the standard deviation. The orange line in (F) and (G) is shown for reference and corresponds to a perfect scaling scenario, where the amount of a cell component increases in direct proportion to cell size so that its concentration does not change. The total amount of protein is very close to perfect scaling, while the amount of GSH increases faster than cell size so that its concentration is higher in larger cells. (I, J) Comparison of GSH concentrations in small, medium, and large cells. Based on the flow cytometry data, 5% smallest, 5% largest and 5% intermediate-sized RPE-1 cells were gated (I), and their GSH concentration (GSH amount per total protein amount) histograms were plotted for each of these size populations (J). The plot shows that the GSH concentration distributions progressively shift towards higher values when cell size increases. The plots in (I-J) are based on the primary data shown in panels (C-D).

While the upregulation of GSH biosynthesis may promote the resistance of larger cells to ferroptosis, such an upregulation alone cannot explain why larger cells become more resistant to ferroptosis induced by the cystine import inhibitor Era2, but not, for example, by the GPX4 inhibitor RSL3 (Chan et al, 2025) (Figs. 2B, S1B). We found previously that upon mTORC1 inhibition cells can evade cystine deprivation-induced ferroptosis by uptake and catabolism of cysteine-rich extracellular proteins, mostly albumin (Armenta et al, 2022) (Fig. S3C). This process involves albumin degradation in lysosomes, predominantly by cathepsin B (CatB), and subsequent export of cystine from lysosomes to fuel the synthesis of glutathione. Large cells undergo proteome rearrangements similar to those occurring upon mTORC1 inhibition (Zatulovskiy et al, 2022). This suggests that large cells may upregulate CatB expression to bypass the Era2-induced cystine import inhibition via system x_c_^-^. To test this hypothesis, we used flow cytometry to measure how the expression of cathepsin B and the system x_c_^-^ cystine/glutamate transporter SLC7A11 (xCT) scales with cell size (Fig. 4B). We found that SLC7A11 concentration modestly decreases, while CatB concentration significantly increases with cell size (Fig. 4B). This shift in the ratio between SLC7A11 and CatB supports the hypothesis that larger cells may rely less on cystine import via system x_c_^-^ and thus become more resistant to system x_c_^-^ inhibition by Era2.

To further test the prediction that larger cells have more GSH to protect against ferroptosis, we measured GSH abundance in different-sized cells using flow cytometry. To do this, we stained the cells with the fluorescent GSH probe monochlorobimane (MCB) (Rice et al, 1986; Shrieve et al, 1988). In addition to MCB, we also stained the same cells with the total protein dye CFSE (carboxyfluorescein diacetate succinimidyl ester) (Lanz et al, 2022; Berenson et al, 2019), which allowed us to calculate relative cell size-dependent changes in glutathione concentrations (amounts of glutathione per unit protein mass; Fig. 4C-H). While total protein amounts scaled in proportion to cell size (Fig. 4G), the amount of MCB-reactive GSH ‘super-scaled’ so that larger cells have higher glutathione concentrations than smaller cells (Fig. 4F,H-J). Overall, these findings indicate that larger cells have higher concentrations of glutathione, likely due to their higher concentrations of GSH producing enzymes and cathepsin B, which could specifically protect them from system x_c_^-^ inhibition-induced ferroptosis.

### Higher concentrations of ACSL4 in smaller cells drives increased membrane lipid peroxidation

Besides the increase in GSH-producing enzyme concentrations with cell size, our proteomics data also indicate that the concentration of another key protein involved in ferroptosis, Acyl-CoA Synthetase Long Chain Family Member 4 (ACSL4), also changed significantly with cell size (Fig. 4A). ACSL4 promotes ferroptosis sensitivity by enriching phospholipid and triacylglycerol pools with ω-6 PUFAs that are prone to peroxidation (Doll et al, 2017) (Fig. 3A). Our proteomics data (Lanz et al, 2022) suggested that ACSL4 concentration decreases with cell size (Fig. 5A). To confirm this finding using flow cytometry, we stained the cells with antibodies against ACSL4. We observed that the ACSL4 concentration indeed decreases with cell size, while the concentration of the control protein β-actin remains constant (Fig. 5B). This suggested the lower lipid peroxidation in larger calls may be due in part to lower ACSL4 abundance.

**Figure 5.**
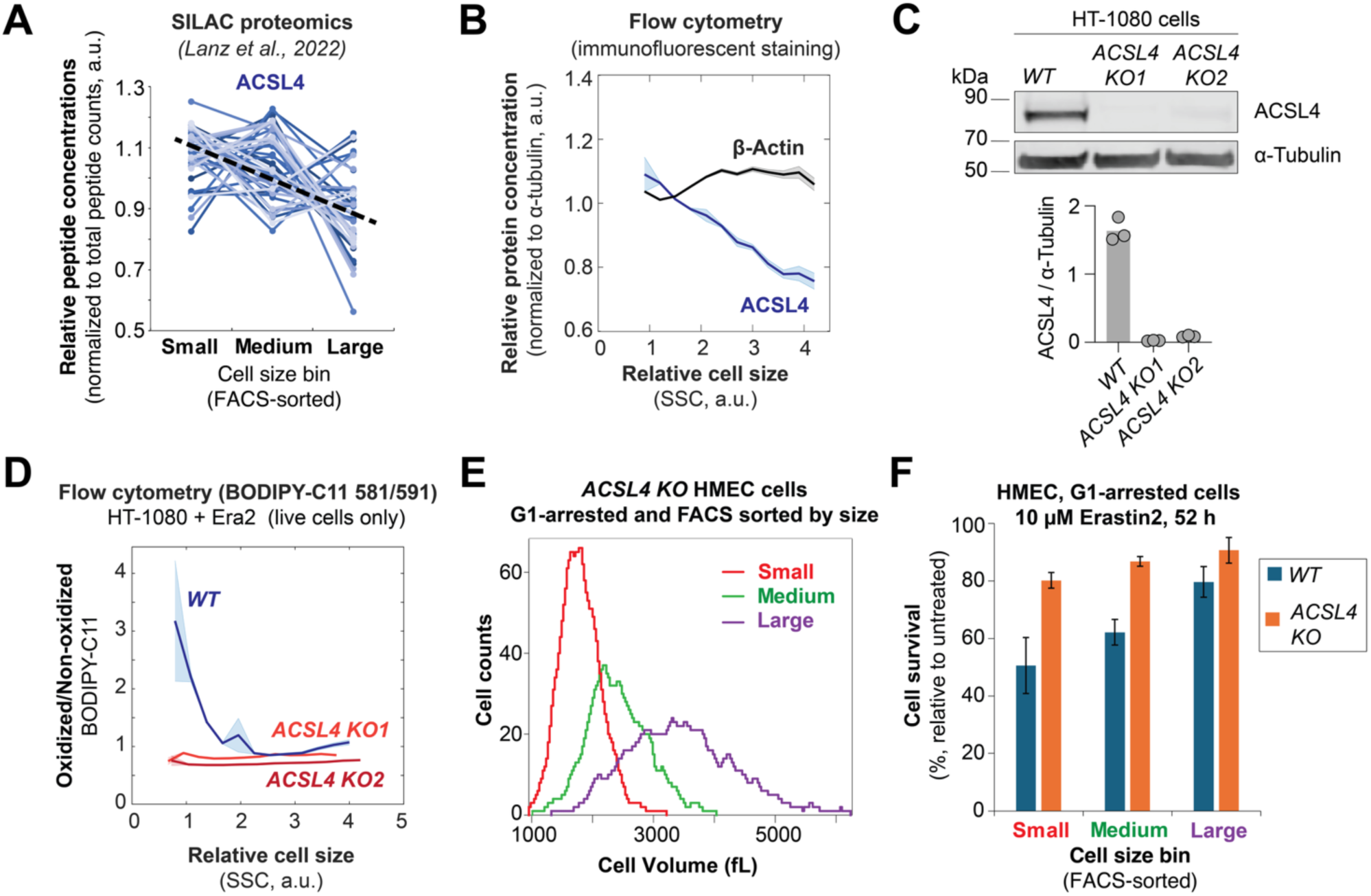
Higher expression of ACSL4 in smaller cells drives increased lipid peroxidation and ferroptosis. (A) Proteomics analysis identified the size-dependent expression of ACSL4, an enzyme that enriches cellular membranes with long polyunsaturated fatty acids prone to peroxidation (Doll et al, 2017). The primary proteomics data were from our previous work (Lanz et al, 2022). Each line in the plot corresponds to a unique peptide from the ACSL4 protein identified by mass spectrometry; the dashed black line indicated the average ACSL4 protein slope. (B) Flow cytometry analysis confirms the decrease of ACSL4 concentration with cell size in HMEC cells. β-Actin was measured as a reference protein, as its concentration does not change with cell size. To calculate the concentrations of ACSL4 and β-Actin, their amounts were measured with flow cytometry using immunofluorescence and were normalized to the amounts of α-Tubulin. The data were binned by cell size, and mean values for each bin were plotted against normalized cell size (solid blue line for ACSL4 and black line for β-Actin). Shaded areas denote the s.e.m. for each bin. (C) Validation of *ACSL4* knockout in HT-1080 cells with immunoblotting. Wild-type (*WT*) HT-1080 cell line and two different *ACSL4* knockout clones (*KO1* and *KO2*) were analyzed using antibodies against ACSL4 and α-Tubulin as a loading control. Bar plot shows the quantification of immunoblotting data from three biological replicates. (D) Deletion of *ACSL4* eliminates the size-dependence of membrane lipid peroxidation in HT-1080 cells. The plot shows the flow cytometry measurements of lipid peroxidation after 16 h 1 µM Era2 treatment in wild-type and *ACSL4 KO* HT-1080 cells (*ACSL4 KO1* and *ACSL4 KO2* are two different knock-out clones) (Magtanong et al, 2019). The ratio between oxidized (green) BODIPY-C11 and non-oxidized (red) BODIPY-C11 fluorescence is plotted as a metric for lipid peroxidation, and the side-scatter parameter (SSC-A) is used as a proxy for cell size. The flow cytometry data were binned by cell size, and mean values of oxidized to non-oxidized BODIPY-C11 ratios were plotted for each bin (blue solid line corresponds to *wild-type* cells, orange and red lines correspond to *ACSL4* gene-disrupted clones *KO1* and *KO2*). Shaded areas denote the s.e.m. for each bin. (E) Cell size distributions of small, medium, and large G1-arrested *ASCL4 KO* HMEC cells isolated by FACS sorting. Prior to FACS sorting, the cells were cultured for 24 h in the presence of 1 µM palbociclib to synchronize cells in G1 phase. Cell size after sorting was measured on a Coulter counter. (F) Cell survival percentage in *WT* and *ACSL4 KO* HMEC cells, sorted into small, medium, and large size bins by FACS. After sorting, the cells were re-plated in the presence of 1 µM palbociclib to keep them in G1 phase. Cells were then treated with 10 µM Era2 for 52 h, and the cell survival percentage was calculated relative to palbociclib-only treated cells. Cell survival percentages in graphs are shown as means ± s.e.m. for n = 3 biological replicates.

To test the role of ACSL4 in size-dependent lipid peroxidation, we measured lipid peroxidation in wild-type and *ACSL4* gene-disrupted HT-1080 cells (Magtanong et al, 2019). We stained the cells with BODIPY-C11 and measured fluorescence with flow cytometry. While lipid peroxidation decreased with increasing cell size in wild-type HT-1080 cells, we did not observe any such size-dependence in *ACSL4 KO* cells (Magtanong et al, 2019) (Fig. 5C,D). This indicated that ACSL4 activity was required for size-dependent changes in basal lipid peroxidation, which likely makes smaller cells more susceptible to ferroptosis.

To test the hypothesis that size-dependent expression of ACSL4 mediates the increased resistance of larger cells to ferroptosis, we FACS-sorted *ACSL4 KO* HMEC cells into small, medium and large size bins and assessed their responses to Era2 (Figs. 5E,F and S4). In this experiment, we synchronized cells in G1 phase using palbociclib prior to cell sorting and also incubated the sorted cells in the presence of palbociclib during Era2 treatment to isolate cell size effects from the previously observed confounding effects of the cell cycle on ferroptosis (Fig. 2B,E). Consistent with our hypothesis, genetic disruption of *ACSL4* reduced the size dependence of the ferroptotic response in Era2-treated HMEC cells (Fig. 5F). We note that while genetic disruption of *ACSL4* significantly reduced the size dependence of ferroptosis, it does not eliminate the size-dependence completely. This might be because ferroptosis is driven by two parallel processes – ACSL4-dependent PUFA incorporation into the membrane and subsequent lipid peroxidation (Fig. 3), both of which depend on cell size.

## DISCUSSION

In this study, we found that larger cells are less susceptible to ferroptotic cell death induced by the system x_c_^-^ inhibitor erastin2 (Fig. 6). This observation may explain previous findings where genetic alterations that reduce cell size also promote ferroptosis, whereas those that increase cell size inhibit it. For instance, deletion of p21, which decreases cell size, sensitizes to ferroptosis, whereas overexpression of p21, increasing cell size, confers ferroptosis resistance (Tarangelo et al, 2018; Venkatesh et al, 2020). Similarly, deletion of the retinoblastoma protein (RB), which also reduces cell size, promotes ferroptosis. Moreover, further deletion of additional RB family members, p107 and p130, in a triple knockout that drastically reduces cell size, increases sensitivity to ferroptosis (Kuganesan et al, 2021; Sage et al, 2000). Here, we demonstrate that isogenic cell populations differing only in size exhibit different susceptibilities to ferroptosis. This implies that many genetic interactions associated with ferroptosis may actually operate indirectly through their effects on cell size.

**Figure 6.**
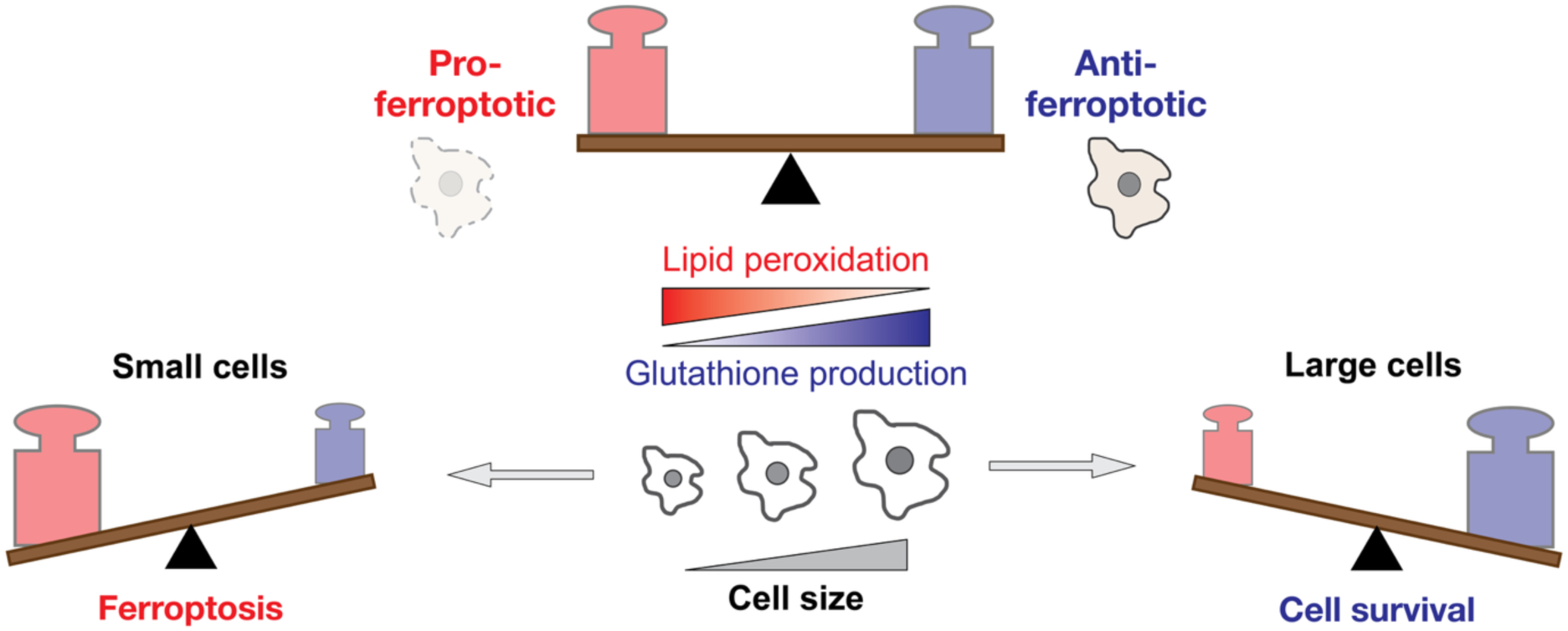
Cell size modulates cell susceptibility to ferroptosis. Larger cells are less prone to Era2-induced cell death because they generate less peroxidized plasma membrane lipids and more glutathione to reduce those toxic peroxidized lipids.

That increased cell size protects against system x_c_^-^ inhibitor-induced ferroptosis has important implications for how this form of cell death interacts with the cell division cycle. Cells in G1 phase of the cell cycle were reported to be more susceptible to ferroptosis (Rodencal et al, 2024; Kuganesan et al, 2023), which suggested that ferroptosis inducers could be used in combination with cancer drugs, like the CDK4/6 inhibitor palbociclib, that arrest cells in G1 phase of the cell cycle (Herrera-Abreu et al, 2024). However, while CDK4/6 inhibitors arrest cells in G1, they do not inhibit cell growth, such that the longer they are arrested, the larger the cells grow (Lanz et al, 2022; Crozier et al, 2023; Manohar et al, 2023). This results in a complex, non-monotonic ferroptotic response dynamics in cells treated with CDK4/6 inhibitors (Fig. 2B,E). Just following CDK4/6 inhibitor treatment, as more and more cells are arrested in G1 phase, cells become more sensitive to both RSL3- and erastin-induced ferroptosis (Kuganesan et al, 2023; Rodencal et al, 2024). However, the longer the cells are arrested, the larger they become, which further promotes their susceptibility to RSL3 (Fig. S1B) but reduces their susceptibility to Era2-induced ferroptosis (Fig. 2B). The fact that the cell cycle arrest and cell size increase have opposing effects on Era2-induced ferroptosis susceptibility could explain why different studies reported seemingly contradictory results, where sometimes an increased and sometimes a decreased or unchanged sensitivity to system x_c_^-^ inhibitors was observed depending on the cell type, duration and type of cell cycle arrest (Lee et al, 2024; Kuganesan et al, 2023; Rodencal et al, 2024). Such complex interplay between the cell cycle and cell size effects on ferroptosis suggests that combination therapies utilizing CDK4/6 inhibitors and ferroptosis inducers would have to carefully choose a dosage schedule. Additionally, our data suggest that previously reported resistance of senescent cells to ferroptosis can at least partially be due to the increased cell size, a well-established hallmark of senescence (Davies et al, 2022).

That small cells are more prone to ferroptosis is likely related to their proteome and metabolome composition differing from that of larger cells. Here, we found that larger cells both generate fewer peroxidized membrane lipids and accumulate more glutathione, which can be used to reduce and thus detoxify the peroxidized lipids (Fig. 6). Likely, these size-dependent changes in lipid oxidation and glutathione abundance are due to differential cell-size-scaling of different metabolic enzymes in the cell. Larger cells also have higher concentrations of iron-chelating protein ferritin and glutathione-producing enzymes GSS and GCLM that protect cells against ferroptosis, and have lower concentrations of the enzyme ACSL4, which promotes ferroptosis. Indeed, the genetic disruption of *ACSL4* greatly reduced the size-dependence of membrane lipid peroxidation, a ferroptosis trigger, and makes ferroptosis less size-dependent. Taken together, these observations indicate that the size-dependent remodeling of the proteome underlies much of the size-dependent sensitivity to ferroptosis.

The fact that larger cell size promotes increased ferroptosis resistance to system x_c_^-^inhibitors (Era2) but, at the same time, decreased resistance to ferroptosis induced by GPX4 inhibitors (RSL3) suggests that the critical roles in these two types of responses are played by different molecular pathways. This finding supports the notion that the ferroptotic cell death response is highly context-dependent and varies dramatically depending on the cell type and class of stimuli inducing ferroptosis (Rodencal et al, 2024; Magtanong et al, 2022; Soula et al, 2020). While a decreased cell sensitivity to Era2 in larger cells is due to size-dependent increase in glutathione production and decrease in unsaturated lipid incorporation into the membrane by ACSL4, the increased sensitivity to RSL3 is reported to be mediated mainly by ER expansion, iron accumulation, and lipid accumulation and remodeling (Chan et al, 2025). We show that large cells may become resistant specifically to Era2 but not RSL3 through the upregulation of lysosomal function, particularly cathepsin B expression, which enables the uptake and catabolism of cysteine-rich extracellular proteins. A size-dependent shift in the ratio between SLC7A11 and cathepsin B makes large cells less dependent on cystine import via system x_c_^-^, and thus, more resistant to Era2. In addition to this, it was reported that RSL3 can induce ferroptosis independently of GPX4 and may target other selenoproteins (DeAngelo et al, 2025; Cheff et al, 2023), which could also contribute to the difference in size-dependent responses to RSL3 and Era2.

In conclusion, our finding that the response of cells to ferroptosis inducers is influenced by cell size point to a broader pattern where cellular decisions are often size-dependent. A general size-dependence of cellular decisions may be driven by widespread changes in protein and metabolite composition taking place as cells grow larger. Indeed, larger cells differ markedly from their smaller counterparts, which likely affects key decisions related to cell death, division, and differentiation. Previous studies, including our own, have explored how cell size influences cell division and contributes to cellular senescence (Lanz et al, 2022; Zatulovskiy et al, 2022; Crozier et al, 2023, 2022; Manohar et al, 2023; Foy et al, 2023; Wilson et al, 2023; Demidenko & Blagosklonny, 2008). Here, we have demonstrated that cell size also plays a significant role in modulating susceptibility specifically to ferroptotic cell death. Furthermore, research in developmental biology shows that cell size can dictate developmental outcomes, as seen in the asymmetrical division of neurosecretory motor neuron neuroblasts in *C. elegans* and cell fate determination in *Arabidopsis* leaves (Sethi et al, 2022; Gong et al, 2023). Thus, we anticipate many cellular decisions will be impacted by cell size, the fundamental mode of cell geometry that sets the basic scale of all intracellular processes.

## ACKNOWLEDGEMENTS

We thank Leslie Magtanong for assistance with the initial cell death screen design and all members of the Zatulovskiy and Skotheim laboratories for valuable discussions and feedback on this project. Cell sorting for this project was performed on an instrument in the Stanford Shared FACS Facility purchased by Parker Institute for Cancer Immunotherapy, and an instrument in the University of Cambridge, Department of Pathology FACS facility. We thank both FACS facilities for their assistance. This work was supported by a Chan Zuckerberg Biohub Investigator Award (J.M.S.), the NIH (P01 CA254867 grant to J.M.S. and R01 GM122923 to S.J.D.), and the Medical Research Council (MR/X020290/1 Career Development Award to E.Z.).

## AUTHOR CONTRIBUTIONS

E.Z. conceived this study. E.Z., M.B.M., S.J.D., and J.M.S. designed the experiments. E.Z., M.B.M., and S.Z. performed the experiments and analyzed the experimental data. E.Z. and J.M.S. wrote the manuscript. E.Z., M.B.M., S.J.D., and J.M.S. edited the manuscript.

## MATERIALS AND METHODS

### Cell culture conditions and cell lines

Non-transformed *hTERT*-immortalized human mammary epithelium cells (HMEC*-hTERT* cells, in this paper referred to simply as HMEC for brevity) were obtained from Stephen Elledge’s laboratory at Harvard Medical School (Solimini et al, 2012) and cultured in MEGM™ Mammary Epithelial Cell Growth Medium (Lonza CC-3150). HT-1080 cells were purchased from ATCC (Manassas, Virginia, USA). Non-transformed *hTERT*-immortalized human retinal pigment epithelium cells (cell line RPE-1) were obtained from the Stearns laboratory at Stanford. Both the HT-1080 and RPE-1 cell lines were grown in Dulbecco’s modification of Eagle’s medium (DMEM) with L-glutamine, 4.5 g/L glucose and sodium pyruvate (Corning), supplemented with 10% FBS (Corning) and 1% penicillin/streptomycin. All cells were cultured at 37°C with 5% CO_2_. HT-1080 cell lines expressing nuclear-localized fluorescent protein mKate2 (Nuc::mKate2) were generated by lentiviral transduction of a viral vector at an M.O.I. of 0.3, that directed the expression of nuclear-localized mKate2 (Forcina et al, 2017). Polyclonal mKate2-expressing populations were selected using puromycin (1.5 mg/mL, 72 h). *ACSL4 KO* HT-1080 and HMEC cells were generated using a CRISPR/Cas9 system, as described previously (Magtanong et al, 2019). RPE-1 cells carrying a genetic construct for a conditional knock-down of cyclin D1 gene (*CCND1)* through doxycycline-inducible *CCND1* shRNA expression were obtained from the Ioannis Sanidas laboratory at Harvard Medical School.

### Immunofluorescence cell staining for flow cytometry

For flow cytometry analysis, cells were grown on dishes to ∼50% confluence and harvested by trypsinization. The cells were then fixed with 3% formaldehyde for 10 min at 37°C and permeabilized with 90% methanol for 30 min on ice. Fixed and permeabilized cells were washed once with PBS, blocked with 3% BSA in PBS for 30 min at 37°C, and then stained with primary antibodies for 2 h at 37°C (rat-anti-alpha-Tubulin (Abcam, #ab6160), mouse-anti-Actin (Sigma-Aldrich, #A2103), rabbit-anti-ACSL4 (Proteintech, #22401-1-AP), rabbit-anti-Cathepsin B (Proteintech, #12216-1-AP), rabbit-anti-xCT/SLC7A11 (Cell Signaling Technology, #12691T)). The cells were then washed twice with a wash buffer (1% BSA in PBS + 0.05% Tween® 20), stained with the fluorophore-conjugated secondary antibodies Alexa Fluor 488 goat anti-rat (Life Technologies, #A11006), Alexa Fluor 594 goat anti-rabbit (Life Technologies, #A11037), and Alexa Fluor 647 goat anti-mouse (Thermo Fisher Scientific, #A32728) at 1:1000 dilution for 1 h at 37°C. The cells were then washed again twice. After this treatment, the cells were resuspended in PBS containing 3 µM DAPI for DNA staining, incubated for 10 min at room temperature, and then analyzed on an Attune NxT flow cytometer (Thermo Fisher Scientific). Around 100.000 cells were typically recorded for each sample, and three biological replicates were performed for each experiment. The flow cytometry gating strategy is shown in Fig. S5. Briefly, single cells were gated based on FSC-A vs SSC-A, then FSC-A vs FSC-H, then SSC-A vs SSC-W plots. From this population of single cells, G1 cells were selected using Hoechst-A vs FSC-A plot (Fig. S5).

For plotting, all protein amounts and cell size values were normalized to the means for each sample. To compensate for the nonspecific background staining of cells with antibodies, we measured the fluorescence of cells stained with nonspecific Isotype Control antibodies. We then performed a linear regression of this nonspecific background signal with cell size and subtracted the background fluorescence corresponding to each cell’s size from the actual anti-ACSL4 fluorescence signal measured for each cell. For all binned flow cytometry data plots, the cells below the 2^nd^ and above the 98^th^ cell size percentiles were excluded to remove the extreme outliers. Then, the remaining data were binned by size and plotted as background-corrected average fluorescence intensity for each bin against the bin’s average cell size. Bins with fewer than 200 cells were excluded from the analysis to reduce noise. For protein concentration calculation, the amount of the indicated protein in the cell was normalized to the amount housekeeping protein alpha-Tubulin, which scales in proportion to cell volume (Lanz et al, 2022).

### Lipid peroxidation measurement with BODIPY-C11 581/591 fluorescent dye

For lipid peroxidation analysis, cells were treated with DMSO or erastin2 (Era2) for 24 h before being harvested by trypsinization. A cell suspension containing ∼200,000 cells was then transferred to a 1.5 mL microfuge tube. Cells were then pelleted by centrifugation (400 x g, 5 min) and resuspended in BODIPY-C11 581/591 (5 μM) dissolved in HBSS. A dead cell stain SYTOX Blue (Molecular Probes, #S34857) was added to cell suspensions at a final concentration of 20 nM to identify and exclude from the analysis all non-intact (dead or dying) cells. Cell suspensions were incubated at 37°C for 20 min and then pelleted (400 x *g*, 5 min) and resuspended in 0.2 mL of PBS buffer. Samples were strained through a cell strainer prior to flow cytometry analysis. Flow cytometry analysis was performed on an Attune NxT flow cytometer (Thermo Fisher Scientific). Oxidation of BODIPY-C11 581/591 was calculated as the ratio of the green fluorescence (BL1 channel, indicates oxidized probe) to the red fluorescence (YL1 channel, indicates unoxidized probe) (Dixon et al, 2012; Pap et al, 1999).

### Single-cell glutathione (GSH) measurements with MCB fluorescent dye

To measure the GSH amounts in different-sized cells using flow cytometry, we stained the cells with the fluorescent GSH probe monochlorobimane (MCB) (Rice et al, 1986; Shrieve et al, 1988). To do this, the cells were harvested by trypsinization, resuspended in PBS at a density of one million cells per mL, and incubated with 40 μM MCB (Invitrogen, #M1381MP). At the same time, we stained with MCB, we also stained the same cells with a total protein dye CFSE (CarboxyFluorescein Succinimidyl Ester). For total protein staining, the CellTrace CFSE dye (Molecular Probes, #C34554) was added to cell suspensions at a 5 μM concentration. The cells were then stained with MCB and CFSE for 20 min in a tissue culture incubator (37°C, 5% CO_2_) in the dark. The reactions were terminated using 1 mL cold complete medium, followed by centrifugation (400 x *g*, 5 min, +4°C). The pelleted cells were then re-suspended in 0.2 mL of PBS, filtered through a 40 µm cell strainer into FACS tubes and placed on ice. The MCB and CFSE fluorescence signals were measured with an Attune NxT flow cytometer (Thermo Fisher Scientific). GSH concentrations (amounts of glutathione per unit protein mass) in individual cells were calculated as a ratio between the MCB signal (glutathione amount) and CFSE signal (total protein amount).

### Intracellular iron detection with fluorescent probes

To measure labile intracellular iron (Fe^2+^) concentration, live RPE-1 cells were stained simultaneously with three fluorescent probes - FerroOrange (Dojindo Laboratories, Japan, # F374-10) to measure the amounts of labile Fe^2+^ in the cell, CellTrace CFSE dye (Molecular Probes, #C34554) to measure total cellular protein amount, and Hoechst 33342 DNA stain (Thermo Scientific, #62249) to determine the cell cycle phase of the cells. Prior to staining, the cells on a dish were washed twice with warm (37°C) serum-free DMEM and harvested by trypsinization, followed by centrifugation and supernatant removal. The cell pellets were then resuspended in a staining solution containing 1 µM FerroOrange, 5 µM CFSE and 20 µM Hoechst in warm serum-free DMEM and incubated at 37°C for 30 min in a 5% CO2 incubator. After the staining the cells were kept on ice and measured on an Attune NxT flow cytometer (Thermo Fisher Scientific). Hoechst signal was used to select G1-phase cells for subsequent analysis.

A similar protocol was used to measure lysosomal Fe^2+^ amounts, but instead of FerroOrange the cells were stained with 1 µM Lyso-FerroRed probe (Dojindo Laboratories, Japan, # L270-10). After Lyso-FerroRed staining, the cells were centrifuged and washed twice with serum-free DMEM prior to the flow cytometry measurement.

### Fluorescence-activated cell sorting (FACS)

Fluorescence-activated cell sorting was used to sort live cells by their size and cell cycle phase, as described previously (Lanz et al, 2022). Cells were harvested from dishes by trypsinization, stained with 20 µM Hoechst 33342 DNA dye in PBS for 15 min at 37°C, and then sorted on a BD FACSAria Fusion flow cytometer. Consecutive SSC-A over FSC-A, and FSC-H over FSC-A gates were used to isolate single cells. Then, G1 cells were gated by DNA content (Hoechst staining). Finally, we collected the 10% smallest and 10% largest cells using the gating based on SSC-A signal as a proxy for cell size. During sorting, all cell samples and collection tubes were kept at 4°C. To determine the cell size distributions of the collected samples, aliquots were taken from each sorted size bin and measured on a Z2 Coulter counter (Beckman). Sorted cells were re-plated for subsequent evaluation of their sensitivity to cell-death-inducing chemicals.

### Microscopy-based analysis of cell death kinetics

For the analysis of cell death susceptibility, FACS-sorted small and large G1-phase cells were seeded on a 384-well plate (100-1500 cells per well), allowed to adhere overnight, and then treated with cell-death-inducing compounds for 72 h. Besides these lethal compounds, the cell culture media also contained 20 nM of the dead-cell dye SYTOX Green (Molecular Probes, #S7020), which permeates dying cells and produces a strong green fluorescent signal in cell nuclei. Cells were imaged on the Essen IncuCyte Zoom and analyzed using scalable time-lapse analysis of cell death kinetics (STACK) as described below. Two or three independent biological replicates were performed for each experimental condition. Each of the biological replicates contained two technical replicates for each condition that were then averaged for cell-death sensitivity quantification. Cell death responses were measured using the scalable time-lapse analysis of cell death kinetics (STACK) technique (Forcina et al, 2017; Inde et al, 2020). Cell lines stably expressing nuclear-localized mKate2 were incubated in medium with 20 nM SYTOX Green and the indicated cell-death-inducing compounds. Counts of live (mKate2+) and dead (SG+) objects were obtained from images acquired every 4 h on the Essen IncuCyte Zoom (Essen BioScience, Ann Arbor, MI). The following image extraction parameter values were used for automated image analysis. For SG+ objects (dead cells): Adaptive Threshold Adjustment 3; Edge Split On; Edge Sensitivity −7; Filter Area min 0 μm^2^, max 750 μm^2^. For mKate2+ objects (live cells): Adaptive Threshold Adjustment 2.5; Edge Split On; Edge Sensitivity −2; Filter Area min 50 μm^2^, maximum 8100 μm^2^; Eccentricity max 0.9. For Overlap objects: Filter area min 50 μm^2^, maximum 8100 μm^2^. Counts were exported to MS Excel (Microsoft Corporation, Redmond, WA) and lethal fraction (LF) scores were computed from mKate2+ and SG+ counts, with the additional step of removing ‘overlap’ double positive counts from live cell counts at each timepoint, as described before (Forcina et al, 2017; Inde et al, 2020). LF scores were exported to Prism 9.0.1 (GraphPad Software, La Jolla, CA) for curve fitting and data plotting.

### Immunoblotting

For immunoblotting, cells were lysed in RIPA lysis buffer on ice. Proteins from lysates were separated on 10% SDS-PAGE gels and transferred to nitrocellulose membranes. Membranes were then blocked with SuperBlock™ (TBS) Blocking Buffer (Thermo Fisher Scientific) and incubated overnight at 4°C with primary antibodies in 3% BSA solution in PBS. The primary antibodies used were: anti-ACSL4 (Sigma-Aldrich, #SAB2701949; 1:2500 dilution) and anti-tubulin (Millipore Sigma, #MS581P1; 1:2000 dilution). The primary antibodies were detected using the fluorescently labeled secondary antibodies IRDye® 680LT Goat anti-Mouse IgG (LI-COR 926-68020) and IRDye® 800CW Goat anti-Rabbit IgG (LI-COR 926-32211). Membranes were imaged on a LI-COR Odyssey CLx and analyzed with LI-COR Image Studio software.

### Data quantification, statistical analysis, and software

Lethal fraction calculations and statistical analyses of cell death were performed using Microsoft Excel 14.6.0 (Microsoft Corporation, USA). Flow cytometry data were analyzed using FlowJo 10.6.1 (FlowJo LLC, USA) and Matlab R2022b (MathWorks, USA). Graphing was performed using Prism 9 (GraphPad Software, La Jolla, CA).

## SUPPLEMENTARY DATA

**Figure S1.**
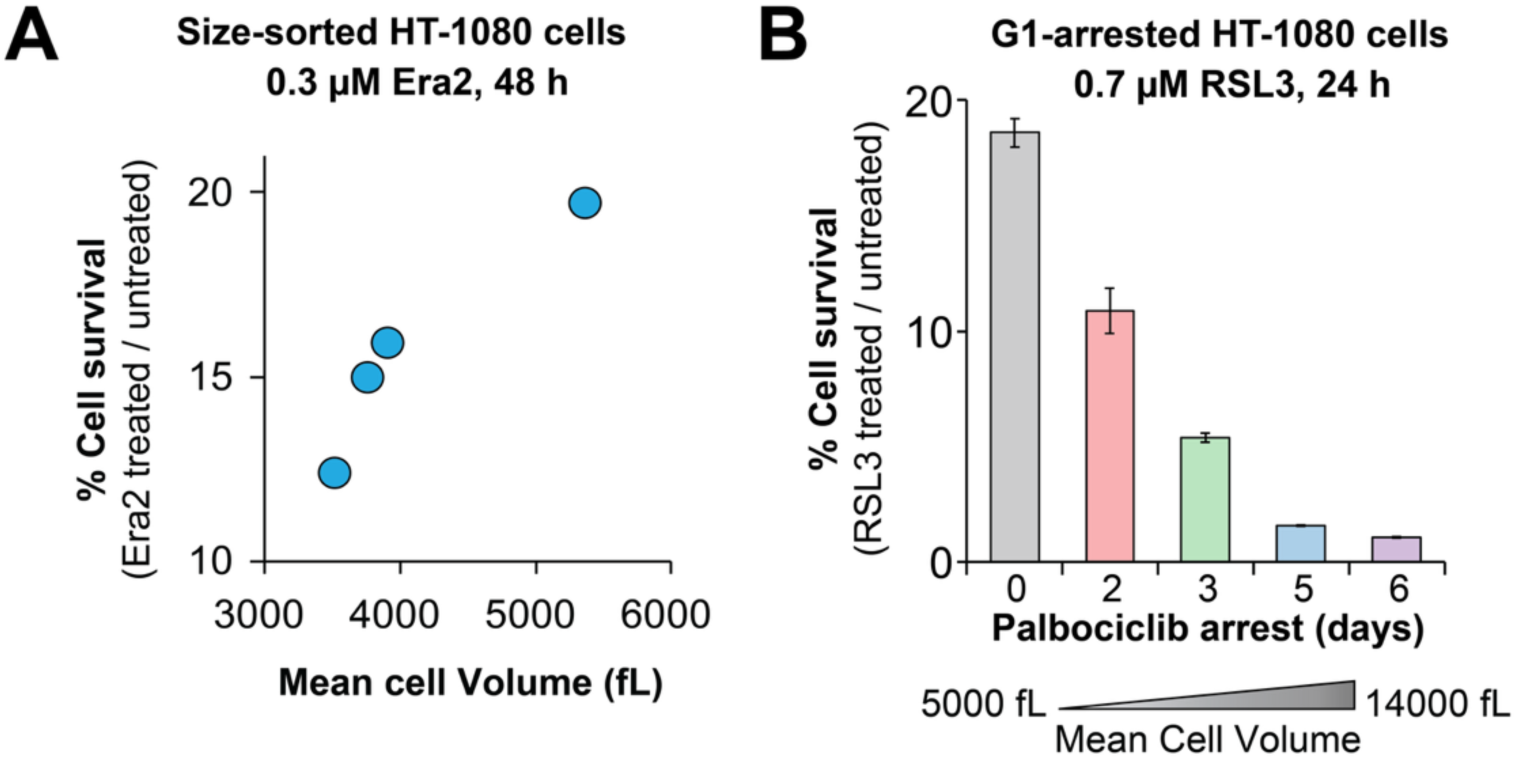
Effects of cell size on Era2- and RSL3-induced ferroptosis susceptibility in HT-1080 cells. (A) Cell survival percentage in HT-1080 cells sorted into four cell size bins using FACS. Cells were treated with 0.3 µM Era2 for 48 h, and cell survival percentage was calculated relative to DMSO-treated cells. The plot shows a representative example from *n* = 2 biological replicates. Each biological replicate included two technical replicates per condition, which were averaged for quantification of cell size and cell death sensitivity. (B) Cell survival percentage in HT-1080 cells whose size was increased through palbociclib-induced cell cycle arrest. After pre-treatment with palbociclib for the indicated numbers of days, the cells were exposed to 0.7 µM RSL3 in the presence of palbociclib for 24 h. Cell survival percentage was calculated relative to the cells treated only with palbociclib. Cell survival percentages are shown as means ± s.e.m.; n = 3 biological replicates.

**Figure S2.**
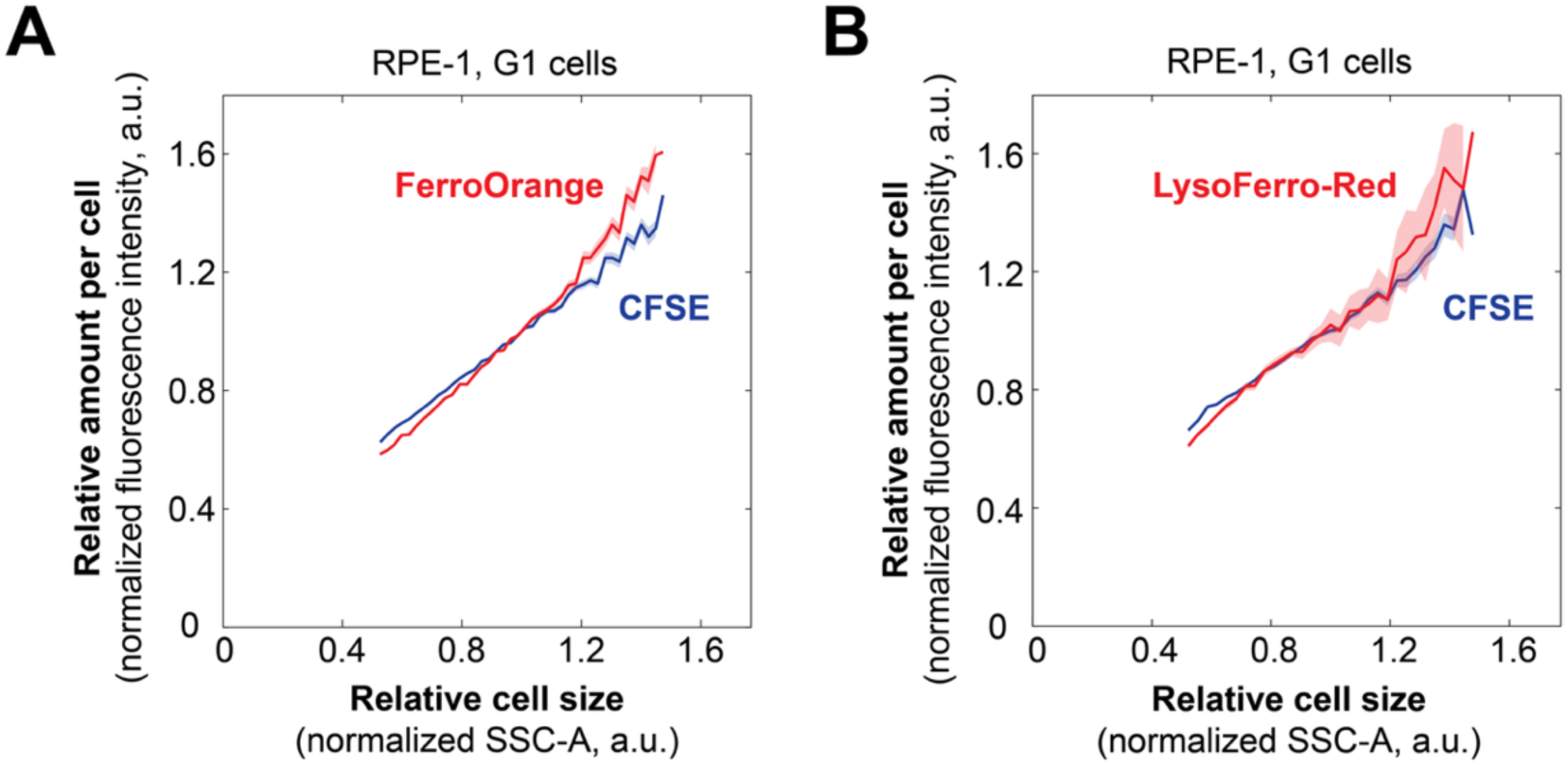
Cell size does not affect the concentrations of labile and lysosome-sequestered iron in RPE-1 cells. (A,B) Flow-cytometry-based measurements of the amount of intracellular labile iron (A) and lysosome-sequestered iron (B) in G1-phase RPE-1 cells. Intracellular labile iron was detected with the FerroOrange fluorescent probe, and lysosomal iron was detected with the Lyso-FerroRed probe. Total cellular protein amount (CFSE staining), which scales in proportion to cell volume, is shown for reference. G1 cells were gated based on Hoechst DNA staining. The data were binned by cell size, and mean values for each bin were plotted against normalized cell size. Shaded areas denote the s.e.m. for each bin. Each sample included 100,000 cells.

**Figure S3.**
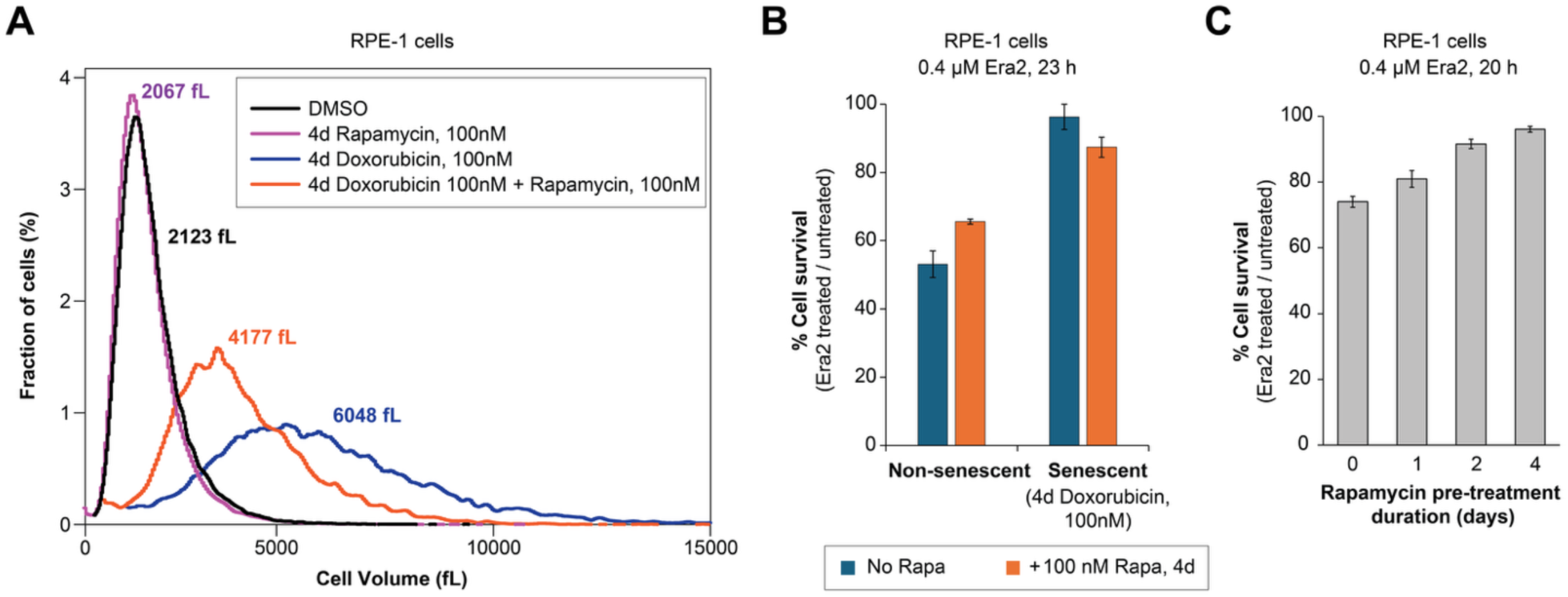
Senescent cells are large and resistant to Era2-induced ferroptosis. (A) Cell size distributions of senescent and non-senescent (asynchronously proliferating) RPE-1 cells, measured with a Coulter counter. The numbers next to the histograms indicate mean cell size ± standard deviation for each condition. 4-day treatment with 100 nM doxorubicin was used to induce senescence in RPE-1 cells. To reduce the size of senescent cells, 100 nM of the mTORC1 inhibitor rapamycin was added to the media during the 4-day doxorubicin treatment. (B) Cell survival percentage in senescent and non-senescent RPE-1 cells treated with 0.4 µM Era2 for 23 h. Senescent cells demonstrate significantly higher resistance to Era2 than non-senescent cells. Senescent cells pre-treated with rapamycin show only a modest reduction in cell survival. Cell survival percentage was calculated relative to the cells under the same conditions treated with DMSO instead of Era2. Error bars show s.e.m., n = 3 biological replicates. (C) Prolonged cell pre-treatment with 100 nM mTORC1 inhibitor rapamycin causes a moderate decrease in cell sensitivity to Era2.

**Figure S4.**
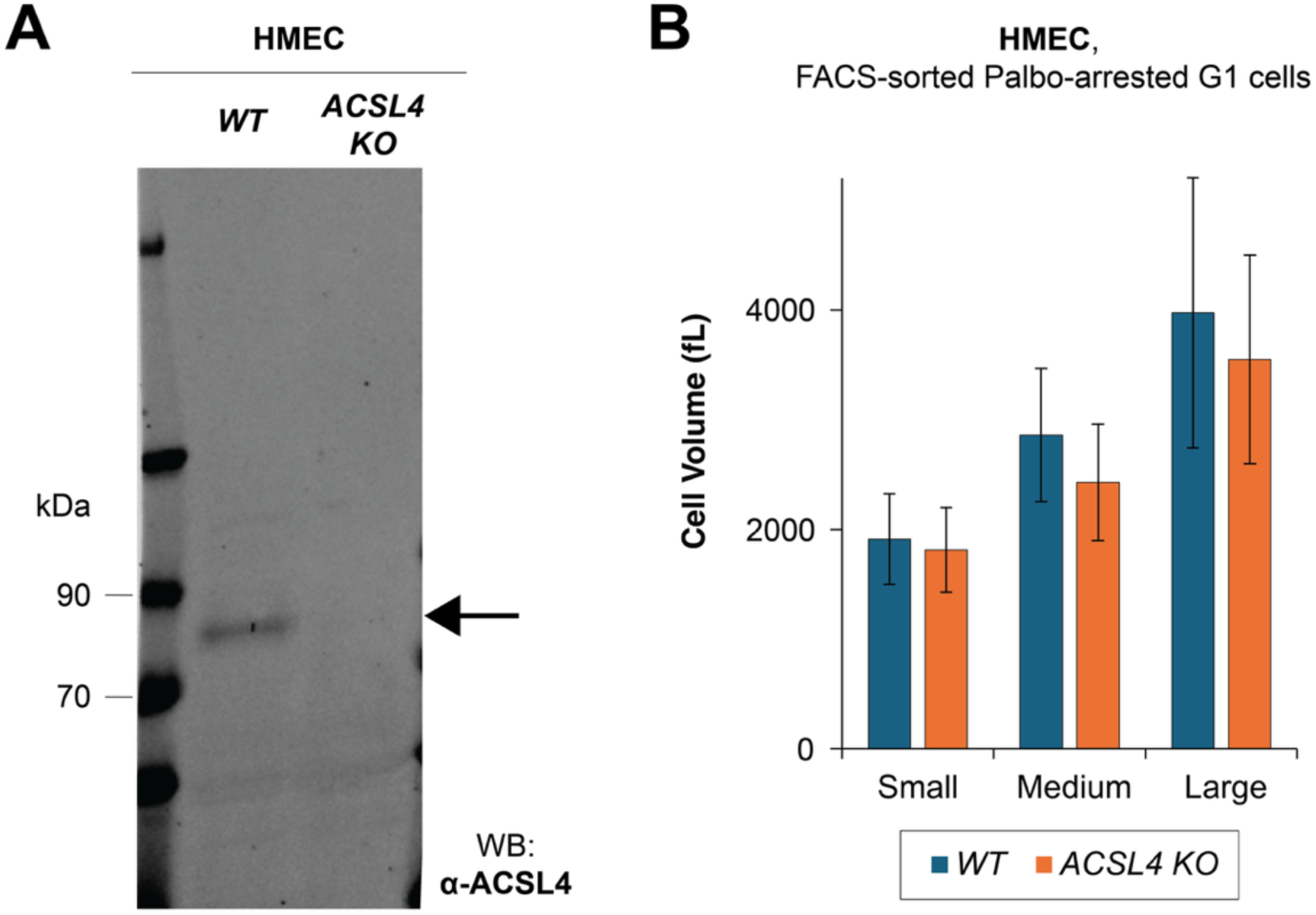
FACS-sorted *ACSL4 KO* HMEC cells have size distributions similar to those of *WT* HMEC cells. (A) Validation of *ACSL4* knockout in HMEC cells with immunoblotting. Wild-type (*WT*) cell line and a mixed (not clonal) population of *ACSL4* knockout cells were analyzed using antibodies against ACSL4. (B) Mean cell sizes of small, medium, and large *WT* and *ACSL KO* HMEC cells after FACS sorting, measured on a Coulter counter. Error bars indicate cell size S.D. Prior to the sorting, the cells were synchronized in G1 phase by a 24 h treatment with 1 µM palbociclib.

**Figure S5.**
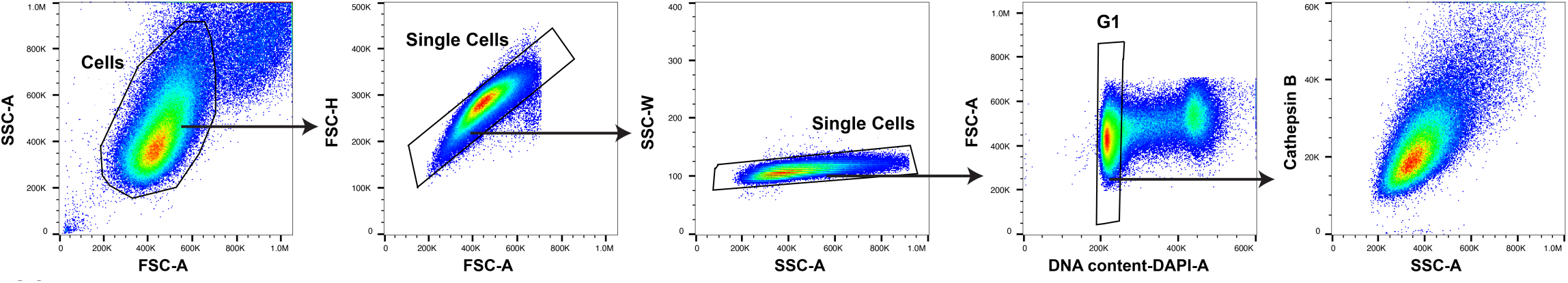
Flow cytometry gating strategy for selecting G1 cells for subsequent scaling analysis. Single cells were gated based on FSC-A vs SSC-A, then FSC-A vs FSC-H, then SSC-A vs SSC-W plots. From this population of single cells, G1 cells were selected using Hoechst-A vs FSC-A plot for subsequent scaling analysis (the example shown is for cathepsin B scaling analysis).

